# New players in the regulatory network of *Vibrio parahaemolyticus* Type VI secretion system 1

**DOI:** 10.1101/464727

**Authors:** Rotem Ben-Yaakov, Dor Salomon

## Abstract

Type VI secretion systems (T6SSs) are widespread, tightly regulated, protein delivery apparatuses used by Gram-negative bacteria to outcompete their neighbors. The pathogen, *Vibrio parahaemolyticus*, encodes two T6SSs. These T6SSs are differentially regulated by external conditions. T6SS1, an antibacterial system predominantly found in pathogenic isolates, requires warm marine-like conditions and surface sensing for activation. The regulatory network that governs this activation is not well understood. In this work, we devised a screening methodology that allows us to easily monitor the outcome of bacterial competitions and thus, to identify mutants that are defective in T6SS1-mediated bacterial killing. The methodology, termed Bacterial Competition Fluorescence (BaCoF), relies on detection of a fluorescent signal as an indicator of the survival and growth of a T6SS-sensitive, GFP-expressing prey that has been co-cultured with mutants derived from a T6SS^+^ attacker of interest. Using BaCoF, we screened a random transposon insertion mutant library and identified genes required for *V. parahaemolyticus* T6SS1 activation, among them TfoY and Tmk. We used epistasis experiments to determine the relationships between the newly identified components and other regulators that were previously described. Thus, we present here a detailed biological understanding of the T6SS1 regulatory network.

## INTRODUCTION

Bacteria use various strategies to carve a niche by outcompeting other bacteria. One of these strategies includes producing antimicrobial proteins that are secreted by toxin-delivery apparatuses, such as the type VI secretion system (T6SS) (Mougous *et al*., 2006; Pukatzki *et al*., 2006; Russell *et al*., 2011). T6SS is a contractile phage tail-like, multi-protein apparatus that is widespread in Gram-negative bacteria (Bingle *et al*., 2008; Boyer *et al*., 2009; Basler *et al*., 2012). It is used to deliver toxins, termed effectors, into neighboring cells in a contact-dependent manner. Effectors can be delivered into either bacterial or eukaryotic neighbors, thus implicating T6SS in both antibacterial competition and virulence (Pukatzki *et al*., 2007; Russell *et al*., 2011). Notably, bacteria encode cognate immunity proteins that protect them against self-intoxication by their own antibacterial T6SS effectors (Russell *et al*., 2011, 2012).

Production of antibacterial arsenals, such as the T6SS, is energy consuming. Therefore, bacteria evolved to regulate these weapons and activate them under specific conditions in which they are required to enhance fitness. Indeed, T6SSs were shown to be tightly regulated in many bacteria, and they may be induced by external conditions and cues such as quorum sensing (Zheng *et al*., 2010), salinity (Salomon *et al*., 2013), temperature (Salomon *et al*., 2013), mucin (Bachmann *et al*., 2015), chitin (Borgeaud *et al*., 2015), surface sensing (Salomon *et al*., 2013), and membrane damage (Basler *et al*., 2013; Ho *et al*., 2013).

*Vibrio parahaemolyticus*, a Gram-negative halophylic bacterium, which is a major cause of seafood-borne gastroenteritis worldwide (Newton *et al*., 2012; Zhang and Orth, 2013) and of acute hepatopancreatic necrosis disease (AHPND) in shrimp (Tran *et al*., 2013; Lai *et al*., 2015), encodes two functional T6SSs (T6SS1 and T6SS2) (Yu *et al*., 2012; Salomon *et al*., 2013). These T6SSs were shown to be differentially regulated by various external cues in the clinical isolate RIMD 2210633. T6SS2 is found in all *V. parahaemolyticus* isolates; however, its role remains unknown. It is active in low salinity (1% NaCl), at low temperatures (23°C), and is repressed upon surface sensing (which can be mimicked in suspension by the polar flagella inhibitor phenamil) (Salomon *et al*., 2013). In contrast, T6SS1, an antibacterial system predominantly found in clinical and AHPND-causing isolates (Yu *et al*., 2012; Li *et al*., 2017), is active under warm marine-like conditions and requires high salinity (3% NaCl), warm temperatures (30°C), and surface sensing for activation (Salomon *et al*., 2013).

We previously described regulators that contribute to the activation of T6SS1 in *V. parahaemolyticus*. Two proteins encoded within the T6SS1 gene cluster (spanning *vp1386-vp1420*), VP1391 and VP1407, act as positive regulators of T6SS1 and are required for its activation (Salomon *et al*., 2013; Salomon *et al*., 2014), whereas the histone nucleoid structuring protein (H-NS) acts as a negative regulator and represses the expression of T6SS1 when the external conditions are not ideal for inducing the system (Salomon *et al*., 2014). OpaR, the high cell density quorum-sensing master regulator, also acts as a negative regulator of this system and its deletion de-represses T6SS1 expression (Salomon *et al*., 2013). However, it remains largely unknown how this bacterium senses and translates surface, salinity, and temperature conditions into a T6SS1-activating signal. Moreover, we do not know whether additional components found outside of the T6SS1 gene cluster are required for its activity (e.g., domesticated core-genome components).

In this study, we set out to identify components that govern the activity of *V. parahaemolyticus* T6SS1. To this end, we devised a fluorescence-based bacterial competition assay that provides a simple, intermediate-throughput method for screening transposon insertion libraries to identify mutants impaired in T6SS-mediated antibacterial activity. Using this method, we identified new components required for T6SS1 activity, and we also determined their role within the T6SS1 regulatory network.

## Results

### Fluorescence-based assay to identify mutants defective in T6SS-mediated antibacterial activity

To identify components that are required for *V. parahaemolyticus* T6SS1 activity, we first sought to develop an assay that would allow us to screen a large number of *V. parahaemolyticus* mutants searching for those impaired in T6SS1-mediated antibacterial activity. To this end, we devised a screening methodology termed <u>B</u>acterial <u>Co</u>mpetition <u>F</u>luorescence (hereafter BaCoF), which uses a green fluorescent protein (GFP) output to detect the outcome of bacterial competitions. GFP serves as an indicator of the survival and growth of a T6SS-sensitive, GFP-expressing prey that has been co-cultured with mutants derived from a T6SS^+^ attacker. Competition with a T6SS^+^ attacker results in prey death and thus no GFP signal, whereas competition with a T6SS-defective mutant allows prey growth and hence, the subsequent detection of the GFP signal.

Briefly, the BaCoF methodology (Fig. 1A) comprises the following steps: i) generating a random transposon insertion mutant attacker library; ii) picking individual attacker mutants into 96-well plates and allowing them to grow overnight; iii) adding T6SS1-sensitive prey that constitutively express GFP in the 96-well plates; iv) spotting the attacker:prey mixtures onto T6SS-inducing, solid media plates; v) following incubation, evaluating the fluorescence of spotted mixtures (detecting fluorescence indicates the lack of prey killing by the mutant attacker, which is thus considered a potential hit); vi) streaking bacterial mixtures containing potential hits onto selective plates to isolate the attacker; and vii) sequencing flanking regions of the transposon to identify the position of its insertion into the genome. The detailed methodology is found in the Methods section.

**Figure 1.**
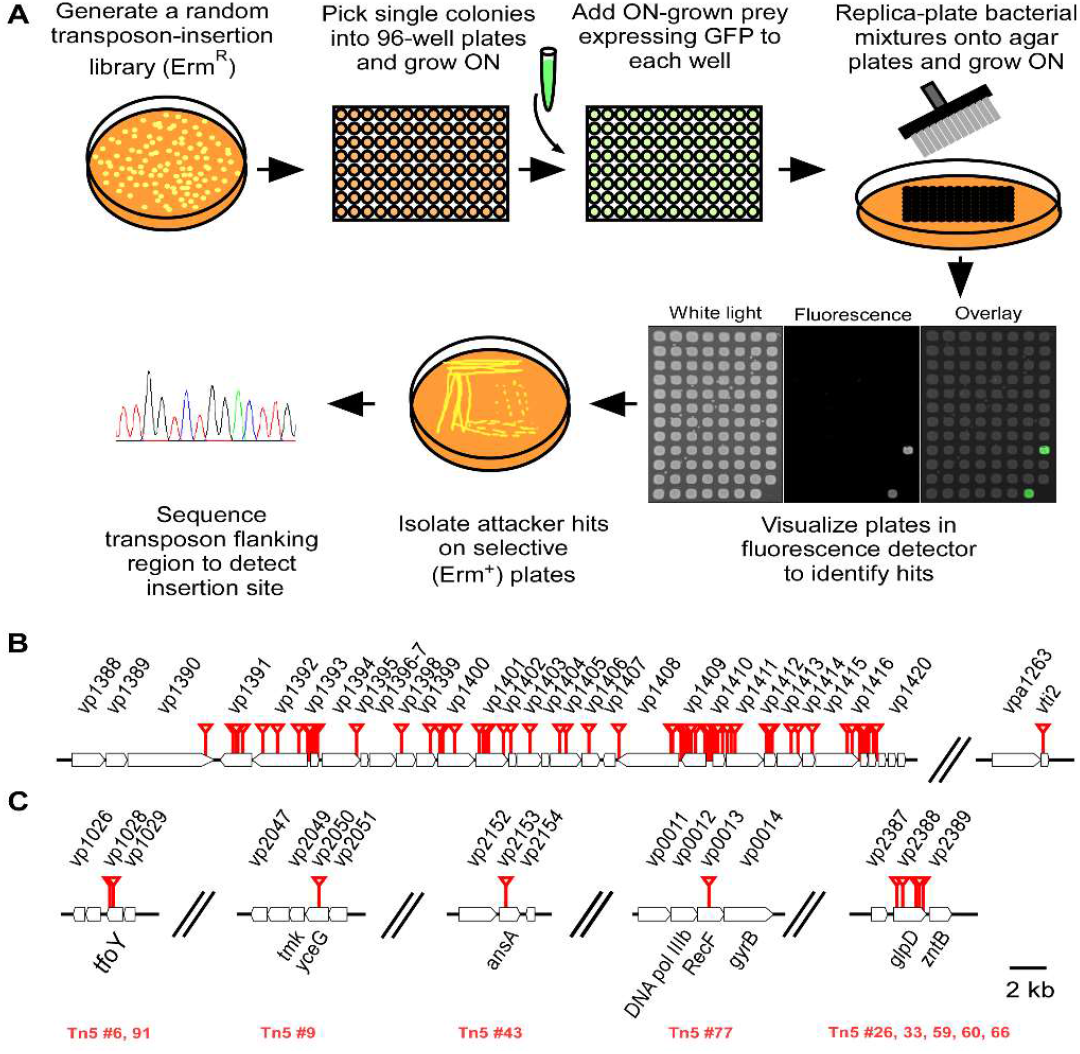
Bacterial Competition Fluorescence (BaCoF) screen reveals *V. parahaemolyticus* mutants defective in T6SS1-mediated bacterial killing. (**A**) Schematic representation of the BaCoF methodology. (**B-C**) Genomic map of BaCoF hits’ transposon insertion sites (red triangles) within known T6SS1 gene clusters and modules (B) and previously non-T6SS1-associated operons (C). Arrows indicate the direction of gene transcription. Locus numbers are listed above and gene names or annotations are listed below. In C, serial numbers of transposon (Tn5) mutant BaCoF hits are shown below in red.

### Applying BaCoF to *V. parahaemolyticus* T6SS1

To apply the BaCoF screen on *V. parahaemolyticus* T6SS1, we used the *V. parahaemolyticus* RIMD 2210633 derivative, POR1, as the parental attacker strain (previously shown to use its T6SS1 to kill competing bacteria under warm marine-like conditions), and the POR1 T6SS1-sensitive derivative, Δ*vp1415-6*, constitutively expressing GFP from a plasmid, as prey. Δ*vp1415-6* harbors a deletion in the T6SS1 effector/immunity pair, *vp1415/6*, and is thus unable to protect itself against attackers that deliver the effector VP1415 via T6SS1 (Salomon *et al*., 2014).

After screening ~12,000 individual mini-Tn5 transposon mutant attackers, we collected 95 potential hits that allowed the growth of the GFP-expressing, T6SS1-sensitive prey. Of these potential hits, 76 had a transposon inserted within the T6SS1 gene cluster (*vp1386-vp1420*) or in a known T6SS1 immunity gene (*vti2*) (Fig. 1B). We expected that mutations within the T6SS1 cluster would affect the antibacterial activity because the cluster encodes core components essential for T6SS1. Therefore, these hits were not further investigated. The remaining 19 potential hits in which the transposon was inserted outside of the T6SS1 gene cluster were subjected to two additional rounds of BaCoF. Ten of the 19 hits passed this validation step and possessed transposon insertion in genes not previously associated with *V. parahaemolyticus* T6SS1: *vp1028*, *vp2050*, *vp0013*, *vp2153*, and *vp2388* (Fig. 1C). These ten potential hits were chosen for further investigation.

### Identification of genes required for T6SS1 induction and activity

First, we determined the ability of the potential hits to mediate T6SS1-dependent bacterial killing by using a quantifiable bacterial competition assay. All ten hits were impaired in their ability to intoxicate the Δ*vp1415-6* prey, although *vp2153*::Tn and *vp0013*::Tn were only partially impaired (Fig. 2A). In addition, since the above self-competition assay only monitors the delivery of a single effector (VP1415), we also measured the bacterial killing ability of these ten hits by monitoring the survival of *E. coli* prey. *E. coli* was chosen as prey, since it was previously shown to be T6SS1 sensitive, and it does not possess immunity genes against the *V. parahaemolyticus* T6SS1 effectors. As shown in Fig. 2B, potential hits with the transposon inserted in *vp1028* and *vp2050* lost their ability to kill *E. coli* prey completely and they were similar to a Δ*hcp1* strain used as a T6SS^-^ control. However, only an intermediate effect on bacterial killing was observed for potential hits with the transposon inserted in *vp0013*, *vp2153*, and *vp2388*.

**Figure 2.**
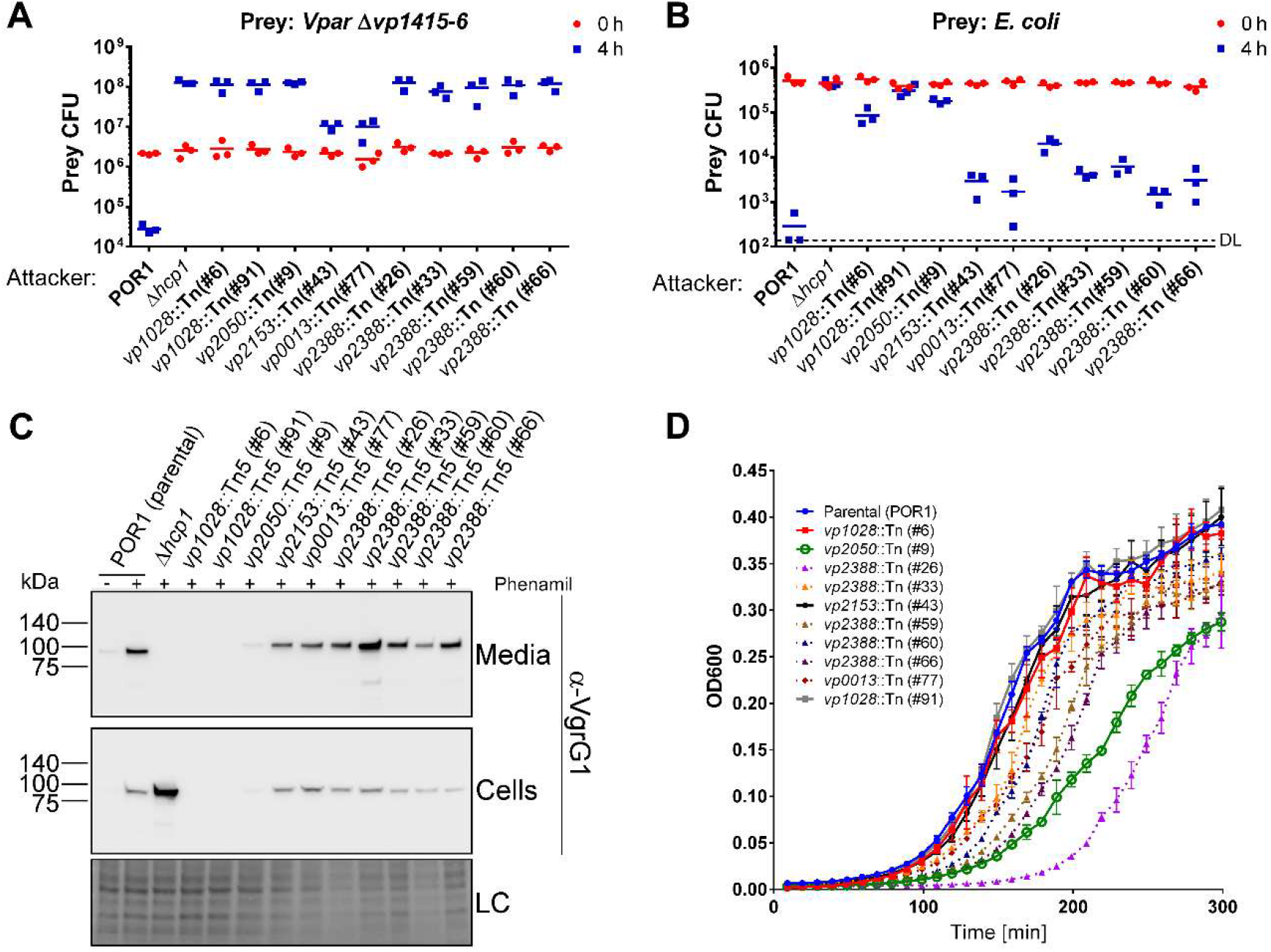
BaCoF hits present T6SS1-mediated bacterial killing, secretion, and growth defects. (**A-B**) Viability counts of *V. parahaemolyticus* POR1/Δ*vp1415-6* (A) and *E. coli* DH5α (B) prey before (0 h) and after (4 h) co-culture with the indicated *V. parahaemolyticus* attackers on MLB media (containing 3% NaCl) at 30°C. (**C**) Expression (cells) and secretion (media) of VgrG1 were detected by immunoblotting using specific antibodies against VgrG1. Loading control (LC), visualized as trihalo compounds’ fluorescence of the immunoblot membrane, is shown for total protein lysates. (**D**) Growth of *V. parahaemolyticus* strains in MLB at 30°C is shown as OD_600_ measurements. Data are mean ± SD, n=3.

Hampered killing activity could have resulted from inability to induce T6SS1 expression following the sensing of external cues (surface, temperature, and salinity), or from inability to properly fire the effector-decorated T6SS tail-tube. To distinguish between these possibilities, we monitored the expression and secretion of the hallmark T6SS-secreted tail-tube component, VgrG1, under T6SS1-inducing conditions. Hits with the transposon inserted in *vp1028* and *vp2050* were unable to express VgrG1 (and thus unable to secrete it). In contrast, hits with the transposon inserted in *vp0013*, *vp2153*, and *vp2388* exhibited VgrG1 expression and secretion levels comparable to the parental POR1 strain (Fig. 2C).

Since hits in *vp0013*, *vp2153*, and *vp2388* exhibited parental-level VgrG1 secretion (Fig. 2C) and were only partially defective in their *E. coli* killing ability (Fig. 2B), we suspected that the effect on bacterial killing could have indirectly resulted from impaired growth of some transposon insertion mutants. Of the mutants lacking bacterial killing activity against *E. coli*, transposon insertions in *vp1028* caused no apparent growth defect. However, insertion in *vp2050* had a dramatic growth defect. Of the mutants retaining intermediate killing activity, only transposon insertion in *vp2153* had no apparent growth defect (Fig. 2D). These results suggest that, whereas the effect of transposon insertions in *vp1028* on bacterial killing did not result from a growth defect, it is possible that the effect of the *vp2050* insertion did. Moreover, it is also plausible that the intermediate killing ability of insertions in *vp0013* and *vp2388* resulted from reduced growth rather than impaired T6SS1 activity *per se*.

Next, we investigated whether the effects on T6SS1 activity can be separated from the effect that some of the hits had on growth. We reasoned that if the defect in T6SS1 activity was only the result of diminished growth or of a general defect in protein secretion across bacterial membranes, then similar effects should be observed in other protein secretion systems. Therefore, we tested how the transposon insertions affected the activity of another protein secretion system, T6SS2, by monitoring the expression and secretion of the hallmark secreted tail-tube component, Hcp2. The hits that were dramatically affected in T6SS1 activity exhibited no defect in Hcp2 expression and secretion, whereas *vp2388*::Tn and *vp0013*::Tn did exhibit reduced Hcp2 secretion levels, compared with the parental strain (Fig. S1). Moreover, we were surprised to find reproducibly high levels of Hcp2 in the secreted fraction of the *vp2050*::Tn mutant, compared with the parental strain. Since T6SS1 induction was nearly completely abolished in this mutant, the finding that T6SS2 is active indicated that the growth defect observed in this mutant was not accompanied by a general secretion defect. Thus, these results suggest that the defect in T6SS1 induction in *vp2050*::Tn might not necessarily be linked to its growth defect.

Taking the above results together, we decided to focus our subsequent efforts on the hits in which T6SS1 activity was abolished (i.e., *vp1028*::Tn and *vp2050*::Tn). The other potential hits that resulted in only an intermediate effect on T6SS1-mediated antibacterial activity were not included in subsequent analyses because we concluded that either: i) we cannot separate their growth defect from a general defect on protein secretion (*vp0013*::Tn and *vp2388*::Tn), or ii) the effect on T6SS1 activity was minor (*vp2153*::Tn).

### TfoY is a positive regulator required and sufficient for T6SS1 activation

Two of our BaCoF hits had a transposon inserted in *vp1028*, which encodes TfoY. Ectopic expression of TfoY from a plasmid restored the bacterial killing ability of the two transposon mutants, indicating that the mutation in *tfoY* caused the loss of T6SS1 activity (Fig. S2). Indeed, deletion of the *tfoY* open reading frame (ORF) recapitulated the phenotype observed for *vp1028*::Tn and abolished T6SS1 antibacterial activity (Fig. 3A,B).

**Figure 3.**
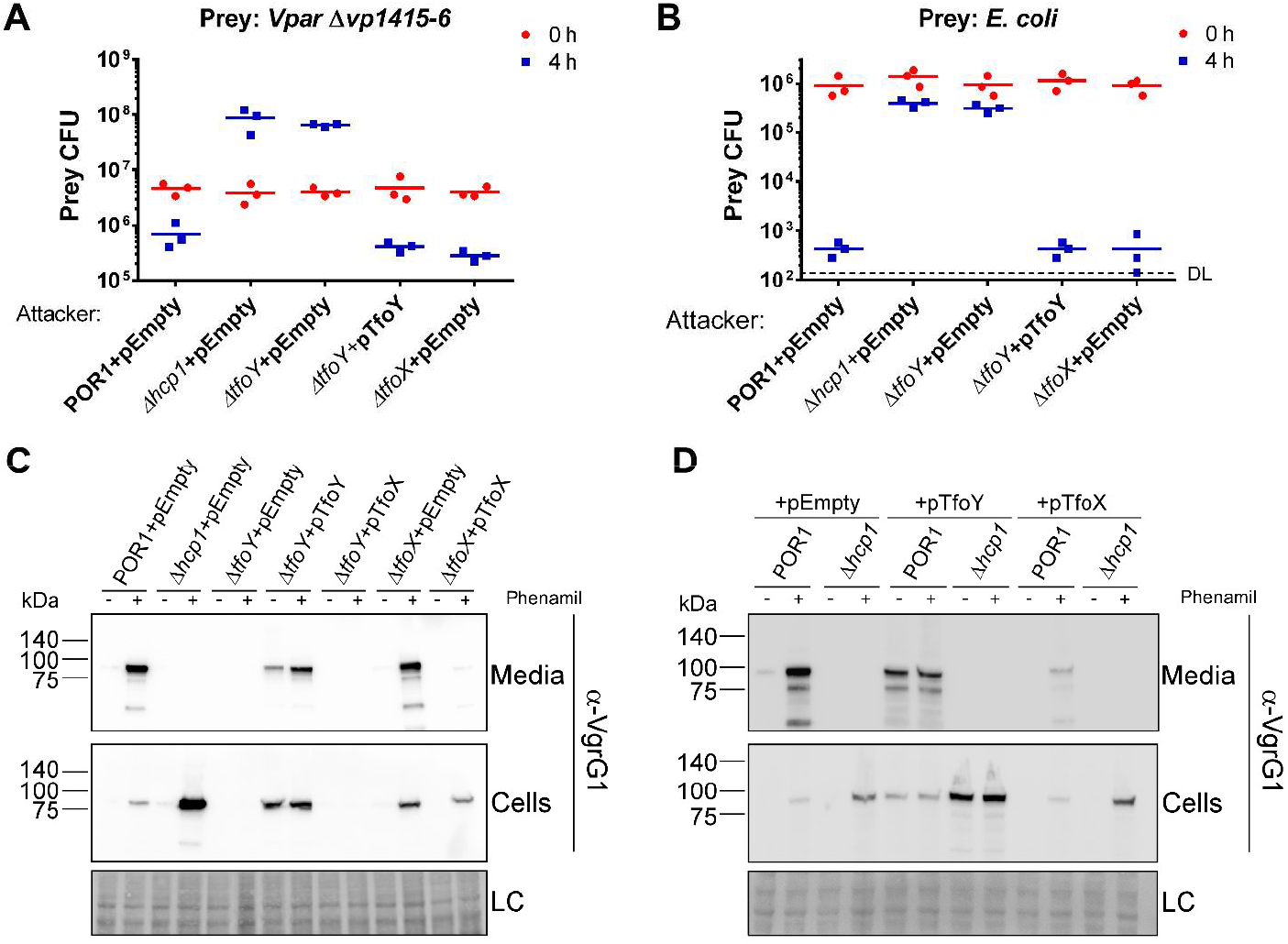
TfoY is an essential positive regulator of *V. parahaemolyticus* T6SS1. (**A-B**) Viability counts of *V. parahaemolyticus* POR1/Δ*vp1415-6* (A) and *E. coli* DH5α (B) prey before (0 h) and after (4 h) co-culture with the indicated *V. parahaemolyticus* attackers on MLB media (containing 3% NaCl) and containing 0.1% L-arabinose at 30°C. (**C-D**) Expression (cells) and secretion (media) of VgrG1 were detected by immunoblotting using specific antibodies against VgrG1. Loading control (LC), visualized as trihalo compounds’ fluorescence of the immunoblot membrane, is shown for total protein lysates. As indicated, the strains contained arabinoseinducible expression vectors of TfoY (pTfoY), TfoX (pTfoX), or an empty vector (pEmpty).

Both TfoY and its paralog, TfoX, were recently shown to orchestrate the activity of T6SS in *V. cholerae*. TfoX was found to be a master regulator of *V. cholerae* T6SS because it was required for chitin-mediated induction of T6SS, and its over-expression induced T6SS induction and antibacterial activity even in the absence of chitin (Borgeaud *et al*., 2015). TfoY was shown to be required for the constitutive activity of *V. cholerae* T6SS in the strain V52, and its deletion resulted in greatly reduced bacterial killing activity and Hcp expression. Moreover, over-expression of TfoY bypassed the requirement for chitin induction of T6SS in strains in which the system is not constitutively active (Metzger *et al*., 2016).

Since *V. cholerae* T6SS differs from *V. parahaemolyticus* T6SS1 in gene content, regulatory components, and activating cues, we investigated whether the concerted regulation of T6SS in *V. cholerae* by both TfoY and TfoX also occurs in *V. parahaemolyticus* T6SS1. We found that whereas TfoY was necessary for surface-induced activation of T6SS1 in *V. parahaemolyticus* (mimicked in suspension by the addition of the polar flagella inhibitor, phenamil (Salomon *et al*., 2013)), TfoX (encoded by *vp1241*) was not required, since T6SS1 activation in Δ*tfoX* resembled that of the parental strain (Fig. 3A-C). Moreover, overexpression of TfoX from a plasmid could not compensate for the loss of T6SS1 activity in Δ*tfoY*, whereas TfoX overexpression had a negative effect on VgrG1 secretion in the presence of TfoY (Fig. 3C,D). We also found that overexpression of TfoY, but not of TfoX, resulted in T6SS1 induction that bypassed the requirement for surface sensing activation (mimicked by the addition of phenamil) (Fig. 3D). Notably, deletion of either *tfoY* or *tfoX* had no effect on bacterial growth (Fig.S3). These results suggest that whereas TfoY is a major positive regulator of surface-induced T6SS1 in *V. parahaemolyticus*, TfoX has a negative effect on the system when it is over-expressed. Thus, the roles of TfoY and TfoX in the T6SS regulatory network differ between *V. parahaemolyticus* and *V. cholerae*.

### Thymidylate kinase is required for T6SS1 activation

The mutant containing a transposon insertion in *vp2050* was severely hampered in its ability to express and secrete VgrG1, as well as in its ability to kill competing bacteria and grow (Fig. 2A-C). Ectopic expression of VP2050 from a plasmid did not rescue these phenotypes, suggesting that the mutation in *vp2050* was not the direct cause of the defect in T6SS1 activation (Fig. 4A,B). This result prompted us to investigate whether the inserted transposon caused a polar effect. Indeed, ectopic expression of the gene found directly downstream of *vp2050*, namely, *vp2049*, rescued the defect in bacterial killing, as well as VgrG1 expression and secretion, and even the growth defect observed in this transposon mutant (Fig. 4). These results indicated that the insult to T6SS1 activation and to growth in this mutant resulted from the polar effect of the transposon insertion on *vp2049*. Ectopic expression of VP2049 even reversed the increased activity of T6SS2 observed in the *vp2050*::Tn mutant (Fig. S4). *vp2049* is an essential gene encoding thymidylate kinase (Tmk), an enzyme that catalyzes the phosphorylation of thymidine 5’-monophosphate (dTMP) to form thymidine 5’-diphosphate (dTDP) in both de novo and salvage pathways of dTTP synthesis (Reynes *et al*., 1996). Therefore, we were unable to generate a Δ*vp2049* strain. Given the annotated activity of VP2049, the mechanism underlying its requirement for T6SS1 activation is not immediately apparent. Notably, it appears that Tmk does not actively play a positive regulatory role in T6SS1 activation because its overexpression did not bypass the need for surface sensing (Fig. S5). Taken together, these results suggest a role for Tmk, or one of the downstream products of its enzymatic activity, in activating T6SS1.

**Figure 4.**
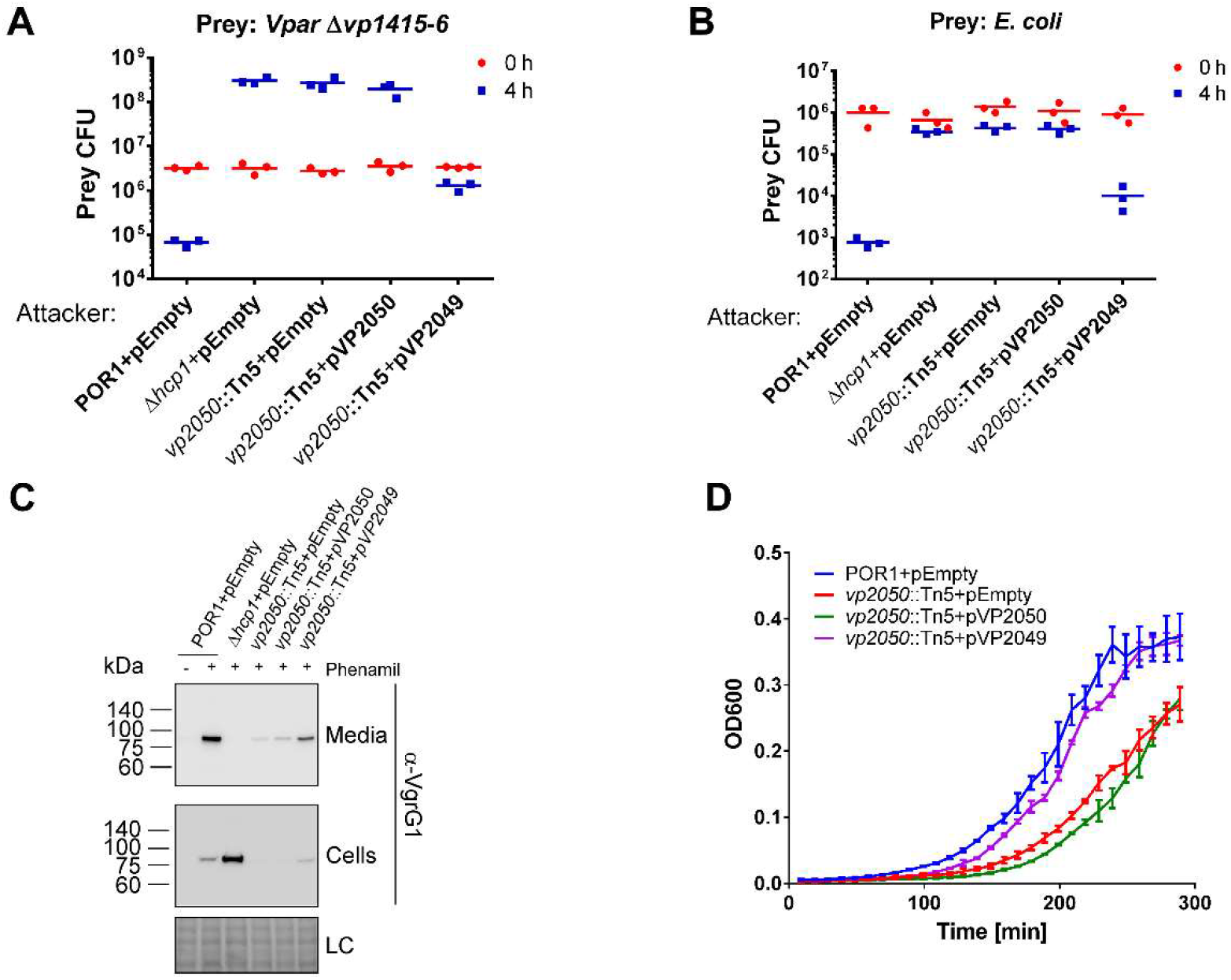
VP2049 (Tmk) is required for T6SS1 activity. (**A-B**) Viability counts of *V. parahaemolyticus* POR1/Δ*vp1415-6* (A) and *E. coli* DH5α (B) prey before (0 h) and after (4 h) co-culture with the indicated *V. parahaemolyticus* attackers on MLB media (containing 3% NaCl) and 0.1% L-arabinose at 30°C. (**C**) Expression (cells) and secretion (media) of VgrG1 were detected by immunoblotting using specific antibodies against VgrG1. Loading control (LC), visualized as trihalo compounds’ fluorescence of the immunoblot membrane, is shown for total protein lysates. (**D**) Growth of *V. parahaemolyticus* strains in MLB containing 0.1% L-arabinose and 250 µg/ml kanamycin at 30°C is shown as OD_600_ measurements. Data are mean ± SD, n=3. As indicated, the strains contained arabinose-inducible expression vectors of VP2050 (pVP2050), VP2049 (pVP2049), or an empty vector (pEmpty).

### A complex regulatory network of negative and positive regulators controls T6SS1 activation

Identifying TfoY as a major positive regulator of *V. parahaemolyticus* T6SS1 prompted us to determine its role and position within the T6SS regulatory network. To this end, we performed epistasis experiments and determined the relationships between known T6SS1 regulators. We constructed mutant strains by combining deletions of T6SS1 negative (*opaR* or *hns*) and positive (*tfoY*, *vp1391*, or *vp1407*) regulators, as well as strains in which positive regulators are overexpressed in the background of a deletion in another positive regulator. We then monitored the effects of these genetic interactions on T6SS1 activity, as measured by antibacterial activity and by VgrG1 expression and secretion. De-repression of T6SS1 by deletion of *hns* required the presence of all three positive regulators (*tfoY*, *vp1391*, and *vp1407*) (Fig. 5A,B). In contrast, whereas de-repression by deletion of *opaR* required *vp1391* and *vp1407*, a double deletion of both *opaR* and *tfoY* (Δ*opaR*/Δ*tfoY*) retained an intermediate activity of T6SS1 (Fig. 5A,C). These findings suggest that H-NS and OpaR repress T6SS1 differently, whereby de-repression by removal of OpaR, but not by removal of H-NS, can bypass the need for activation via TfoY. Moreover, TfoY-mediated activation of T6SS1 was dependent on the presence of both *vp1391*, and *vp1407* (Fig. 5D,E). Notably, neither the single- nor the double-gene deletion strains that were tested exhibited any growth defect, compared to the parental strain (Fig. S6). Taken together, these epistasis interactions reveal a complex regulatory mechanism governing the activation of *V. parahaemolyticus* T6SS1.

**Figure 5.**
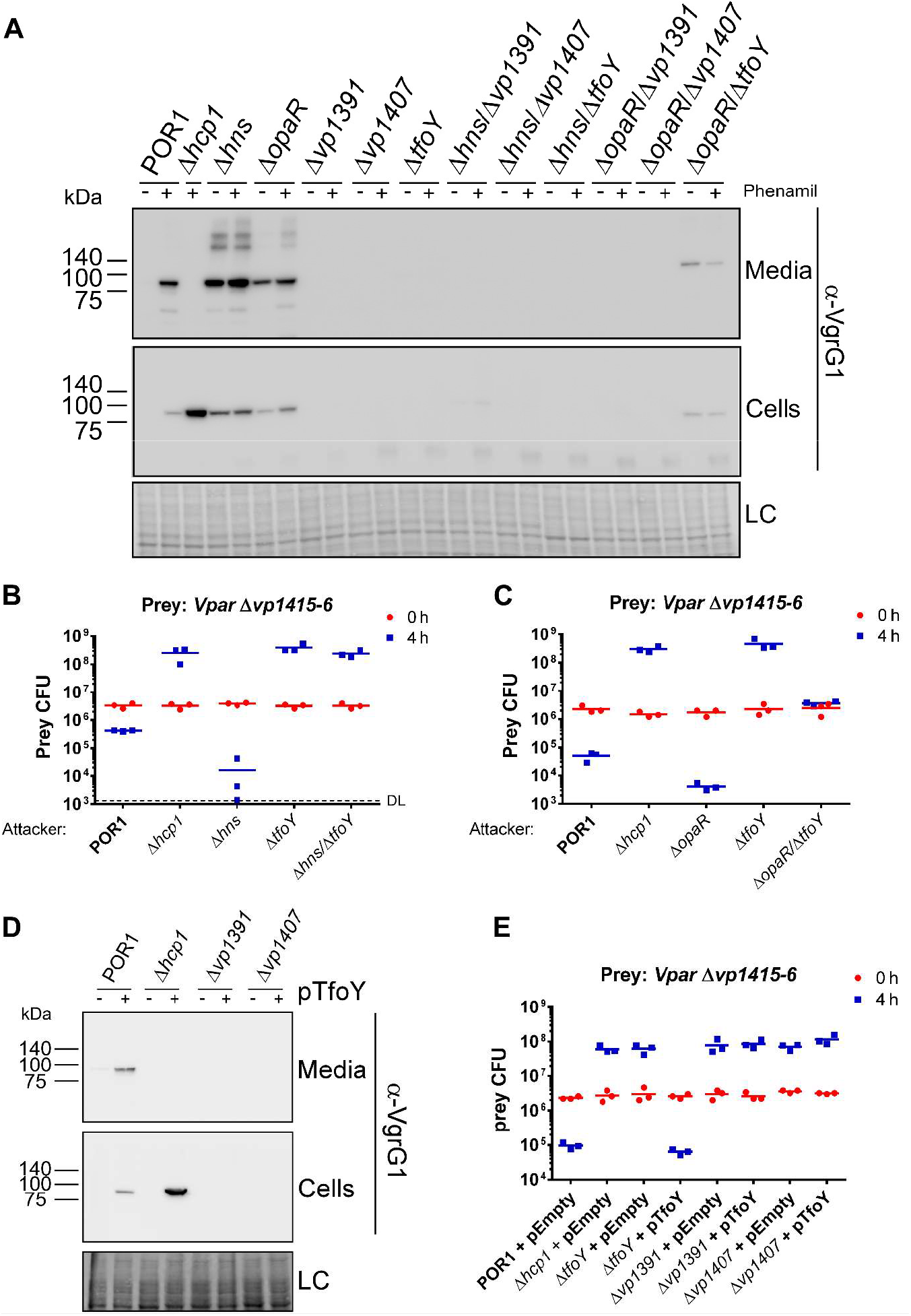
T6SS1 is regulated by a complex network of positive and negative regulators. (**A**) Expression (cells) and secretion (media) of VgrG1 were detected by immunoblotting using specific antibodies against VgrG1. Loading control (LC), visualized as trihalo compounds’ fluorescence of the immunoblot membrane, is shown for total protein lysates. (**B-C**) Viability counts of *V. parahaemolyticus* POR1/Δ*vp1415-6* prey before (0 h) and after (4 h) co-culture with the indicated *V. parahaemolyticus* attackers on MLB media (containing 3% NaCl) at 30°C. (**D**) Expression (cells) and secretion (media) of VgrG1 were detected by immunoblotting using specific antibodies against VgrG1. Loading control (LC), visualized as trihalo compounds’ fluorescence of the immunoblot membrane, is shown for total protein lysates. (**E**) Viability counts of *V. parahaemolyticus* POR1/Δ*vp1415-6* prey before (0 h) and after (4 h) co-culture with the indicated *V. parahaemolyticus* attackers on MLB media containing 0.1% L-arabinose at 30°C. As indicated, the strains contained the arabinose-inducible expression vector of TfoY (pTfoY) or an empty vector (pEmpty).

## Discussion

In this work, we devised a methodology, termed BaCoF, which can be used to screen large numbers of bacterial mutants in order to identify those in which antibacterial activity has been modified. BaCoF relies on fluorescence output as an indicator of bacterial competition outcomes. Using BaCoF, we screened *V. parahaemolyticus* transposon mutants and identified new players in the T6SS1 regulatory network.

Most of the *V. parahaemolyticus* transposon mutants that lost their ability to mediate T6SS1-dependent bacterial killing had a transposon inserted within the T6SS1 gene cluster (*vp1386*-*vp1420*). This result was to be expected, since the T6SS1 gene cluster encodes all the T6SS core components known to be required for normal activity (Boyer *et al*., 2009; Salomon *et al*., 2013). Whereas most genes in the T6SS1 cluster were represented at least once in the list of T6SS1-defective mutants identified, there were a few that were not, such as *vp1386*-*vp1389*, *vp1395*-*vp1396-7* (we recently showed that the region previously thought to encode two proteins, VP1396 and VP1397, actually contains only a single open reading frame termed *vp1396-7*. The erroneous annotation resulted from a mistake in the original deposited sequence (Li *et al*., 2017)), and *vp1418*-*vp1420*. This implies that these unrepresented genes, which are not known T6SS core components (Boyer *et al*., 2009), are not essential for T6SS1 functionality. Indeed, our previous work showed that *vp1388* and *vp1389* encode a T6SS1 effector/immunity pair, and that they are not required for T6SS1 activity (Salomon *et al*., 2014). Moreover, *vp1417*-*vp1420* appear to be diverged duplications of the immunity gene *vp1416*, which is required to provide immunity against the T6SS1 effector VP1415 (Salomon *et al*., 2014). Therefore, it is possible that *vp1418*-*vp1420* are not required to provide immunity against self-intoxication by VP1415, but rather, are used to provide immunity against divergent effectors from other *Vibrio* strains, which would make them expendable under the conditions of our BaCoF assay.

Two transposon insertion hits in our BaCoF assay revealed that *vp1028*, known as *tfoY*, is a required positive regulator of T6SS1. Previous work identified the TfoY homolog in *V. cholerae* as a positive regulator of T6SS. However, since the cues that activate the T6SSs, as do the T6SSs themselves, differ between *V. choerae* and *V. parahaemolyticus*, we continued to investigate the role of TfoY in the regulation of *V. parahaemolyticus* T6SS1. Indeed, we found that the effects of TfoY and of its paralog, TfoX, are quite different in these two vibrios. Whereas TfoY played a positive role in both T6SSs, TfoX did not seem to play a positive role in the regulation of *V. parahaemolyticus* T6SS1. On the contrary, TfoX overexpression resulted in diminished levels of T6SS1 activity. This negative effect of TfoX on *V. parahaemolyticus* T6SS1 can be explained by a previous observation made by Borgeaud *et al*., who found that overexpression of TfoX in *V. cholerae* resulted in down-regulation of TfoY (Borgeaud *et al*., 2015), which is evidently essential for proper activation of *V. parahaemolyticus* T6SS1.

Our finding that overexpression of TfoY bypasses the need for surface sensing induction of T6SS1 suggests that under natural conditions surface sensing activates TfoY, which in turn, activates *V. parahaemolyticus* T6SS1. A similar observation was made with TfoX and the T6SS inducer chitin in *V. cholerae* (Borgeaud *et al*., 2015). Thus, it seems that the major positive activator, as well as the extracellular cue, differ between *V. cholerae* T6SS and *V. parahaemolyticus* T6SS1. Nevertheless, it is interesting that the paralogs TfoY and TfoX play a major role in activating antibacterial T6SSs in these two *Vibrio* species. Since both TfoX and TfoY are highly conserved in vibrios (Pollack-Berti *et al*., 2010), it is plausible that they also control the activity of other *Vibrio* T6SSs. Interestingly, even though TfoY is a central positive regulator of *V. parahaemolyticus* T6SS1, which is induced by surface sensing, apparently it can be bypassed and the system can also be activated by a different pathway. A double deletion of *tfoY* and the negative regulator *opaR* (Δ*tfoY*/Δ*opaR*) exhibited intermediate T6SS1 activity. Thus, it is possible that under certain conditions, quorum sensing regulation (via OpaR) can control T6SS1 activity independent of surface sensing and TfoY.

Using epistasis experiments, we further demonstrated the role of TfoY and of previously identified positive and negative regulators in the T6SS1 regulatory network. As summarized in Fig. 6, we propose a model in which, upon sensing the T6SS1-inducing conditions and cues (surface sensing, 3% NaCl, and 30°C), TfoY is activated and consequently activates the two positive regulators encoded within the T6SS1 gene cluster, VP1391 and VP1407. These two positive regulators are essential for the activation of T6SS1, since they are required for TfoY-mediated activation and also for de-repression of the system, which occurs upon removal of the negative regulation of H-NS or OpaR. The repression mediated by H-NS and OpaR, however, differs, since de-repression by removal of H-NS requires TfoY, whereas that is not the case when OpaR is removed. Therefore, H-NS probably exerts its repression upstream of TfoY, whereas OpaR acts downstream of TfoY to repress T6SS1 activity. This result also implies the existence of a quorum-sensing-dependent, surface sensing/TfoY-independent pathway for T6SS1 activation. In support of this model, OpaR was previously suggested to directly bind to promoter regions of *V. parahaemolyticus* T6SS1 operons, including the promoter of the operon encoding the positive downstream regulator, VP1407 (Zhang *et al*., 2017).

**Figure 6.**
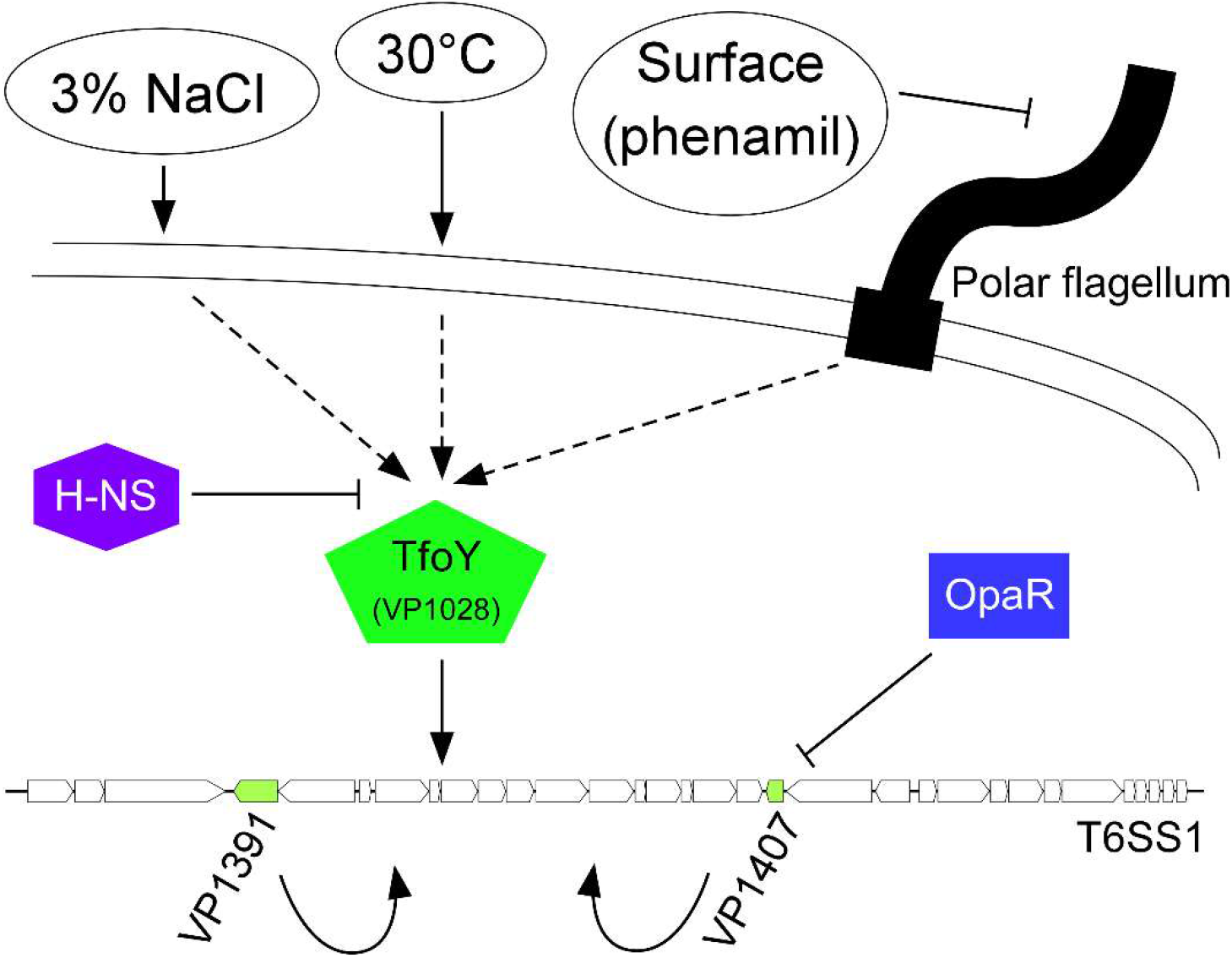
Model of the *V. parahaemolyticus* T6SS1 regulatory network. T6SS1 is induced under warm marine-like conditions and upon surface sensing (via inhibition of the polar flagella). This leads to activation of TfoY, which in turn, activates VP1391 and VP1407 (positive regulators encoded within the T6SS1 cluster) that turn on the antibacterial T6SS1. The quorum-sensing master regulator, OpaR, represses T6SS1 downstream of TfoY, possibly by direct binding to promoters of T6SS1 cluster operons, whereas H-NS represses the system upstream of TfoY.

Note that a previous study also implicated ToxR and AphA in the regulation of *V. parahaemolyticus* T6SS1 (Zhang *et al*., 2017); however, we did not include them in our epistasis analyses since we could not detect any effect of these regulators on T6SS1 activity in our previous work (i.e., their deletion had no negative effect on T6SS1-mediated secretion or antibacterial activity) (Salomon *et al*., 2014). The discrepancy between our previous results and those of Zhang *et al*. could stem from the fact that experiments performed by Zhang *et al*. were conducted under T6SS1 non-inducing conditions (i.e., in the absence of surface sensing activation and at 37°C; conditions under which we previously demonstrated that T6SS1 is inactive (Salomon *et al*., 2013; Salomon *et al*., 2014)). In contrast, our previous work, as well as the current one, examined the effect of regulators under T6SS1-inducing conditions.

Another BaCoF hit that had a dramatic effect on T6SS1 activity was *vp2049*, which encodes Tmk. Although the transposon mutant also exhibited a growth defect, we were able to distinguish the effect on growth from the effect on protein secretion by showing that the negative effect on T6SS1 activation was not evident for T6SS2. Whereas Tmk was required for proper T6SS1 activation, it does not seem to play an active, positive role in activation, since its overexpression did not bypass the need for surface-sensing-mediated induction. The link between Tmk (or the products of its activity) and T6SS1 activation remains unknown, and we will investigate it further in future studies.

Finally, the BaCoF methodology, used here to identify positive regulators and components of *V. parahaemolyticus* T6SS1, can be easily adapted to other systems. To use BaCoF, one needs a system in which conditions that activate an antibacterial T6SS (or any other antibacterial system) are known, and where a suitable sensitive prey that can grow under these conditions is available and can be fluorescently tagged. BaCoF can also be used to identify negative regulators of an antibacterial system, by searching for mutants in which the antibacterial system becomes active under otherwise non-inducing conditions. We are currently applying BaCoF to map regulatory networks of other *Vibrio* T6SSs.

## EXPERIMENTAL PROCEDURES

### Bacterial strains and media

*Vibrio parahaemolyticus* RIMD 2210633 derivative POR1 (Δ*tdhAS*) (Park *et al*., 2004), used here as the parental strain, and its derivatives, were routinely grown in Marine Lysogeny broth (MLB; LB broth supplemented with NaCl to a final concentration of 3% w/v) or on Marine Minimal Media (MMM) agar (2% w/v NaCl, 0.4% w/v galactose, 5 mM MgSO_4_, 5 mM K2SO_4_, 77 mM K2HPO_4_, 35 mM KH2PO_4_, 20 mM NH_4_Cl, 1.5 % w/v agar) at 30°C. To induce the expression of genes from a plasmid, 0.1% (w/v) L‐arabinose was included in the media. *Escherichia coli* strain DH5α was used for plasmid maintenance and amplification, and as prey in competition assays (see below). *E. coli* strain S17‐1 (λ *pir*) was used for maintenance of pDM4 plasmids and mating (see below), and DH5α (λ *pir*) was used for maintenance of isolated transposon-containing genomic regions (see below). *E. coli* was routinely grown in 2xYT broth (1.6% w/v tryptone, 1% w/v yeast extract, 0.5% w/v NaCl) at 37°C. When necessary, media were supplemented with 30 μg/ml (for *E. coli*) or 250 μg/ml (for *V. parahaemolyticus*) kanamycin, and 10 μg/ml chloramphenicol.

### Plasmids

For arabinose-inducible expression in bacteria, the coding sequences of TfoY (*vp1028*) and TfoX (*vp1241*) were amplified from *V. parahaemolyticus* RIMD 2210633 genomic DNA and inserted into the multiple cloning site of the pBAD/Myc–His vector (Invitrogen) harboring a kanamycin-resistance cassette (Salomon *et al*., 2013) in-frame with the C-terminal *Myc*-6xHis tag (producing pTfoY and pTfoX, respectively). pEVS104 conjugative helper and pEVS170 mini-Tn5 delivery vector were a generous gift from Prof. Eric V. Stabb (Lyell *et al*., 2008).

### Construction of deletion strains

For in-frame deletions of *tfoY* (*vp1028*) and *tfoX* (*vp1241*), 1-kb sequences directly upstream and downstream of each gene were cloned into pDM4, a Cm^R^OriR6K suicide plasmid (O’Toole *et al*., 1996). These pDM4 constructs were inserted into *V. parahaemolyticus* via conjugation by S17-1(λ *pir*) *E. coli*. Transconjugants were selected on MMM agar plates containing chloramphenicol. The resulting transconjugants were plated onto MMM agar plates containing 15% (w/v) sucrose for counter-selection and loss of the *sacB*-containing pDM4. Deletions were confirmed by PCR. The generation of in-frame deletions of *hcp1* (*vp1393*), *vp1391, vp1407, hns* (*vp1133*), and *opaR* (*vp2516*) were described previously (Salomon *et al*., 2013; Salomon *et al*., 2014).

### Bacterial Competition Fluorescence (BaCoF) screen

A mini-Tn5 transposon containing OriR6K and erythromycin resistance (Erm^R^) found on the pEVS170 mini-Tn5 delivery vector was conjugated into the *V. parahaemolyticus* POR1 parental attacker. Transconjugates containing a chromosomally integrated transposon were selected on MMM plates containing Erm (100 µg/ml). Single insertion mutant colonies, as well as the parental POR1 (T6SS1^+^) and the POR1/Δ*hcp1* derivative (T6SS1^-^) (Salomon *et al*., 2013), were picked into 96-well plates containing 100 µl MLB per well. Plates were incubated overnight at 30°C. The following day, 40 µl of overnight-grown T6SS1-sensitive prey *V. parahaemolyticus* POR1/Δ*vp1415-6* (Salomon et al., 2014) containing pGFP (a high copy number plasmid that is stably maintained without selection and constitutively expresses green fluorescent protein (GFP) from the lac promoter (Ritchie *et al*., 2012)) were added to each well. Attacker:prey mixtures were spotted onto MLB agar plates using a stainless steel pin replicator, and plates were left open for 10 minutes to dry. The MLB agar plates were then incubated overnight at 30°C and visualized the following morning using a Fusion FX6 imaging system (Vilber) equipped with a RGB excitation module and a GFP emission filter to identify GFP-positive spots. When spots containing GFP fluorescence were identified (indicating the survival and growth of the T6SS1-sensitive prey), 5 µl of the attacker:prey mixtures from the well in 96-well plate that corresponded to the GFP-positive spots were spread on MMM plates containing Erm (100 µg/ml) to isolate the insertion mutant attackers. The location at which the transposon was inserted into these hits was determined as previously described (Lyell *et al*., 2008). Briefly, chromosomal DNA was isolated from insertion mutant hits, digested with HhaI (which does not cut within the transposon), and then was self-ligated. Ligations were transformed into *E. coli* DH5α (λ *pir*) and selected on LB agar plates containing Erm (100 µg/ml). Circular transposons containing the flanking chromosomal DNA were isolated, and transposon chromosomal DNA junctions were sequenced using the M13 forward primer.

### Bacterial competition assays

Quantitative bacterial competition assays were performed as previously described (Salomon *et al*., 2013). Prey strains, either *E. coli* DH5α or *V. parahaemolyticus* derivatives, harbored a pBAD33 plasmid to provide selectable resistance against chloramphenicol. Assays were repeated at least three times with similar results. Results from a representative experiment are shown.

### Secretion assays

*V. parahaemolyticus* secretion assays for T6SS1 and T6SS2 were performed as previously described (Salomon *et al*., 2013). Briefly, for T6SS1 secretion, 5 ml bacterial cultures at an initial OD_600_=0.18 in MLB media were incubated for 5 h at 30°C in the presence or absence of 20 µM phenamil. For T6SS2 secretion, 5 ml bacterial cultures at an initial OD_600_=0.9 in LB media were incubated for 5 h at 23°C. For maintaining plasmids, appropriate antibiotics were added to the media. To induce expression from pBAD vectors, 0.1% (w/v) L-arabinose was added to the media.

For expression fractions (cells) 1.0 OD_600_ units of cells were collected and re-suspended in 100 μl of Tris-glycine SDS sample buffer x2 (Novex, Life Sciences). Supernatants of volumes equivalent to 10 OD_600_ units were filtered (0.22 µm) and precipitated with deoxycholate and trichloroacetic acid (Bensadoun and Weinstein, 1976). Precipitated proteins were washed twice with ice-cold acetone prior to re-suspension in 20 μl of 10 mM Tris–HCl pH=8.0, followed by the addition of 20 μl of 2x protein sample buffer. Expression and secretion samples were resolved on TGX stain-free gels (Bio-Rad), transferred onto PVDF or nitrocellulose membranes, and immunoblotted with custom-made α-VgrG1 (for T6SS1) or α-Hcp2 (for T6SS2) polyclonal antibodies (Li *et al*., 2017). Loading of total protein lysates was visualized by analysis of trihalo compounds’ fluorescence of the immunoblot membrane. Experiments were performed at least three times with similar results.

### Bacterial growth

Overnight-grown cultures of *V. parahaemolyticus* were normalized to an OD_600_=0.01 in MLB media and transferred to 96-well plates (200 µl per well). For each experiment, n=3 or n=6, as indicated. Cultures were grown at 30°C in a BioTek EPOCH2 or SYNERGY H1 microplate reader with continuous shaking at 205 cpm. OD_600_ readings were acquired every 10 minutes. When the cultures contained pBAD expression vectors, kanamycin (250 µg/ml) and L-arabinose (0.1% w/v) were added. Experiments were performed at least three times with similar results.

## ACKNOWLEDGMENTS

We thank Nika Schwartz and Sacha Audey for technical assistance, and other members of the Salomon lab for helpful discussions. We also thank Oren Kobiler for naming BaCoF.

